# RNA-induced allosteric coupling drives viral capsid assembly in bacteriophage MS2

**DOI:** 10.1101/2023.06.05.543665

**Authors:** Sean Hamilton, Tushar Modi, Petr Šulc, Banu Ozkan

## Abstract

Understanding the mechanisms by which single-stranded RNA viruses regulate capsid assembly around their RNA genomes has become increasingly important for the development of both antiviral treatments and drug delivery systems. Here, we investigate the effects of RNA-induced allostery in a single-stranded RNA virus — *Levivirus* bacteriophage MS2 — using the computational methods of the Dynamic Flexibility Index (DFI) and the Dynamic Coupling Index (DCI). We show that asymmetric binding of RNA to a symmetric MS2 coat protein dimer increases the flexibility of the distant FG-loop and induces a conformational change to an asymmetric dimer that is essential for proper capsid formation. We also show that a point mutation W82R in the FG-loop creates an assembly-deficient dimer in which RNA-binding has no significant effect on FG-loop flexibility. Lastly, we show that the highly flexible disordered FG-loop of the RNA bound asymmetric dimer not only becomes the controller of the rigid FG-loop but enhances its dynamic coupling with all the distal positions in the dimer. This strong dynamic coupling allows highly regulated communication and unidirectional signal transduction that drives the formation of the experimentally observed capsid intermediates.

**Author summary:** The final stage of an RNA virus’ life cycle is the assembly of a protein shell encapsulating the viral genome prior to release from the host organism. Despite rapid advancements in both experimental and theoretical biology since the mid-20th century, little is still known about the underlying mechanisms of viral capsid assembly. However, understanding the biophysical principles of viral capsid assembly would bring us one step closer to developing new biotechnologies such as antivirals that inhibit this critical stage of the life cycle or artificial capsids for targeted drug/vaccine delivery. Although we limit the present study to one simple RNA virus that infects bacteria, we propose that the physical implications can extend to other RNA viruses including the human coronavirus SARS-CoV-2. We also propose that the allosteric regulation by specific protein-RNA interactions might be a general mechanism exploited by many other ribonucleoprotein complexes, such as CRISPR-Cas9, spliceosome or ribosome.

## Introduction

Single-stranded RNA (ssRNA) viruses package their genetic material into a protein capsid as part of their replication cycle [1]. Understanding the mechanism of the packaging has importance for development of antiviral therapeutics, as well as for repurposing for delivery applications in bionanotechnology. The main experimental source of information about viral packaging are the atomistic structures resolved through cryoEM or X-ray crystallography, which however only provide static picture of the assembled virus. Several prior studies included coarse-grained simulation of capsid proteins aggregating around negatively charged ssRNA [2]. The pathways of assembly differ between viruses, and the proposed mechanisms include aggregation around viral genome followed by gradual rearrangement, and sequential growth where upon a capsid binding to the viral genome, the subsequent growth is driven by interactions between domains of capsid proteins. A mechanism of RNA packaging signals has been proposed for MS2 and GA virions, where RNA binding promotes protein-protein interactions between capsids [3].

Understanding the mechanisms of protein allostery is an ongoing major challenge in studies of proteins and their functions and interactions [4]. The RNA-induced allosteric regulation in proteins has so far not received much attention. The growing number of theoretical and experimental studies however point to its role in capsid assembly of several viruses. We use here previously developed methods of Dynamic Flexibility Index (DFI) and the Dynamic Coupling Index (DCI) to study the allosteric regulation of MS2 viral capsids induced by binding of its RNA genome. The DFI quantifies flexibility of respective protein residues by quantifying their displacement under an applied force using linear pertubation theory. DFI has been previously shown to identify residues important to protein function, as mutation introduced at rigid positions result in dyfunctional proteins [5, 6]. The DCI measures coupling between pairs of residues upon perturbation of one of them. Prior studies showd that high DCI socre between distant residues indicated long-range allosteric communication [5, 6]. The details about calculating DCI and DFI and their extension to protein-RNA complexes is provided in Methods.

Our results of application of DCI and DFI to MS2 bacteriophage indicate a mechanism of packaging signals-induces sequential assembly of viral capsids, and, more generally, can also be applied to other RNA-protein complexes for identification of possible RNA-mediated allosteric regulation of proteins.

Researchers have extensively studied the *Levivirus* bacteriophage MS2, a positive-sense ssRNA virus that encodes only four proteins: maturation, lysis, replicase, and coat [7–9]. X-ray crystallography (PDB ID: 1ZDH [10]) has revealed that the MS2 genome is encapsulated by 180 coat proteins that form three quasi-equivalent conformations arranged as 60 asymmetric A/B dimers and 30 symmetric C/C dimers. These dimers create a T=3 icosahedral shell with spherically symmetric three-fold (pseudo-six-fold) and five-fold axes. The FG-loops (residues 66–82) of the symmetric and asymmetric dimers, which make crucial inter-dimer contacts at the three-fold axis (FG-loops A and C) and at the five-fold axis (FG-loops B), have significant structural differences [10, 11].

Upon binding to a 19-nt hairpin operator within the replicase gene, the β-sheet interface of the MS2 coat protein dimer acts as a translational repressor [12–14]. NMR experiments have shown that the coat protein dimer favors the symmetric structure in the absence of genomic RNA, whereas it favors the asymmetric structure when bound to the RNA-hairpin operator [15]. Self-assembly kinetics measurements indicate that the assembly process largely depends on the relative concentrations of coat protein and cognate RNA, with an asymmetric dimer bound to the hairpin operator acting as the nucleation site for capsid assembly [12, 13, 16, 17]. The MS2 coat protein can bind to various non-operator RNA-hairpins with high affinity, suggesting that the genome contains at least 59 other nonsequence-specific hairpins that are necessary for successful assembly [13, 18, 19]. Mass spectrometry has revealed multiple protein-RNA contacts within the fully assembled capsid, providing direct evidence that MS2 viral assembly is mediated via non-covalent interactions at RNA-binding regions referred to as ”packaging signals” [20–23]. Additionally, capsid assembly can be disrupted independently of repressor activity due to single point mutations in the FG-loop [14, 24, 25].

Allosteric control, that is the ability of one part of a protein to control another part through changes in conformation upon ligand binding at the controller site allowing for the integration of multiple signals and the coordination of complex biological processes [5, 6, 26, 27], plays a vital role in capsid assembly [28]. In particular, the conformational change of the dimeric FG-loops, critical to capsid assembly is modulated by RNA binding. In the absence of RNA, the two FG-loops in the unbound dimer are symmetrically coupled. However, RNA binding induces disorder in one of the FG-loops leading to an asymmetric structure.

While ligand-induced allostery is well-understood as a common mechanism for the regulation of enzymatic activity and binding interactions in proteins, allosteric control and its mechanistic insights have not yet been generalized to the interactions between proteins and nucleic acids. Here we investigated the mechanism of RNA induced allostery in virus capsid assembly using our conformational dynamics-based analysis [5, 6]. For allostery to effectively regulate a protein, there must be one domain that controls a separate, usually distant, domain. Our results show that, in the symmetric MS2 coat protein dimer, the two FG-loops act as highly flexible and highly coupled domains that communicate symmetrically. When the 19-nt hairpin RNA binds to its native site on one side of the β-sheet interface, the flexibility of the opposite FG-loop increases, and the symmetric coupling is broken. This induces a cascade that signals for the FG-loop to shift into the rigid conformational state found in the asymmetric dimer. The highly flexible disordered FG-loop of the RNA bound asymmetric dimer becomes the controller of the rigid FG-loop, demonstrating the concept of allostery. This controller-controlled dynamic allows for highly regulated communication and unidirectional signal transduction.

The cause-and-effect nature of RNA-binding also gives a mechanistic insight into capsid assembly. The asymmetric dimers then associate with nearby symmetric dimers to form higher order symmetric complexes seen in the three-fold and five-fold ring intermediates. Upon RNA binding, one FG-loop becomes rigid and acts as a hinge, initiating capsid assembly at the five-fold ring. The flexible FG-loop of the asymmetric dimer acts as a signal transducer that interacts with the flexible FG-loop of a symmetric dimer, promoting cooperativity and long-range communication. This communication creates an allosteric network in which the flexible FG-loop modulates the orientation of an incoming dimer, triggering a cascade that allows each newly associated dimer to become the controller of the growing complex.

## Results and Discussion

### DFI analysis of the MS2 wildtype coat protein dimer

The RNA-binding pathway depicted in Fig 1A drives the transition between two quasi-equivalent conformers of the wildtype MS2 coat protein dimer. In the native state, an RNA hairpin binds to one side of the dimer’s β-sheet interface near FG-loop C but distant from FG-loop D. Binding of RNA triggers a conformational change in FG-loop D, resulting in an asymmetric A/B dimer. The symmetric dimer’s DFI profile before and after RNA-binding is presented in Fig 1B. Without RNA, both FG-loops exhibit equal flexibility, reflecting the dimer’s symmetry. However, immediately after RNA-binding, the two FG-loops’ DFI symmetry is disrupted as the transition to the asymmetric dimer occurs. As the average distance between FG-loop D and the RNA-binding sites is 34.1 Å(calculated in PyMOL), the increased dynamic flexibility of FG-loop D (Fig 1C) indicates the presence of RNA-induced allostery. This observation is consistent with a prior all-atom normal-mode analysis of the same system [29, 30].

**Fig 1.**
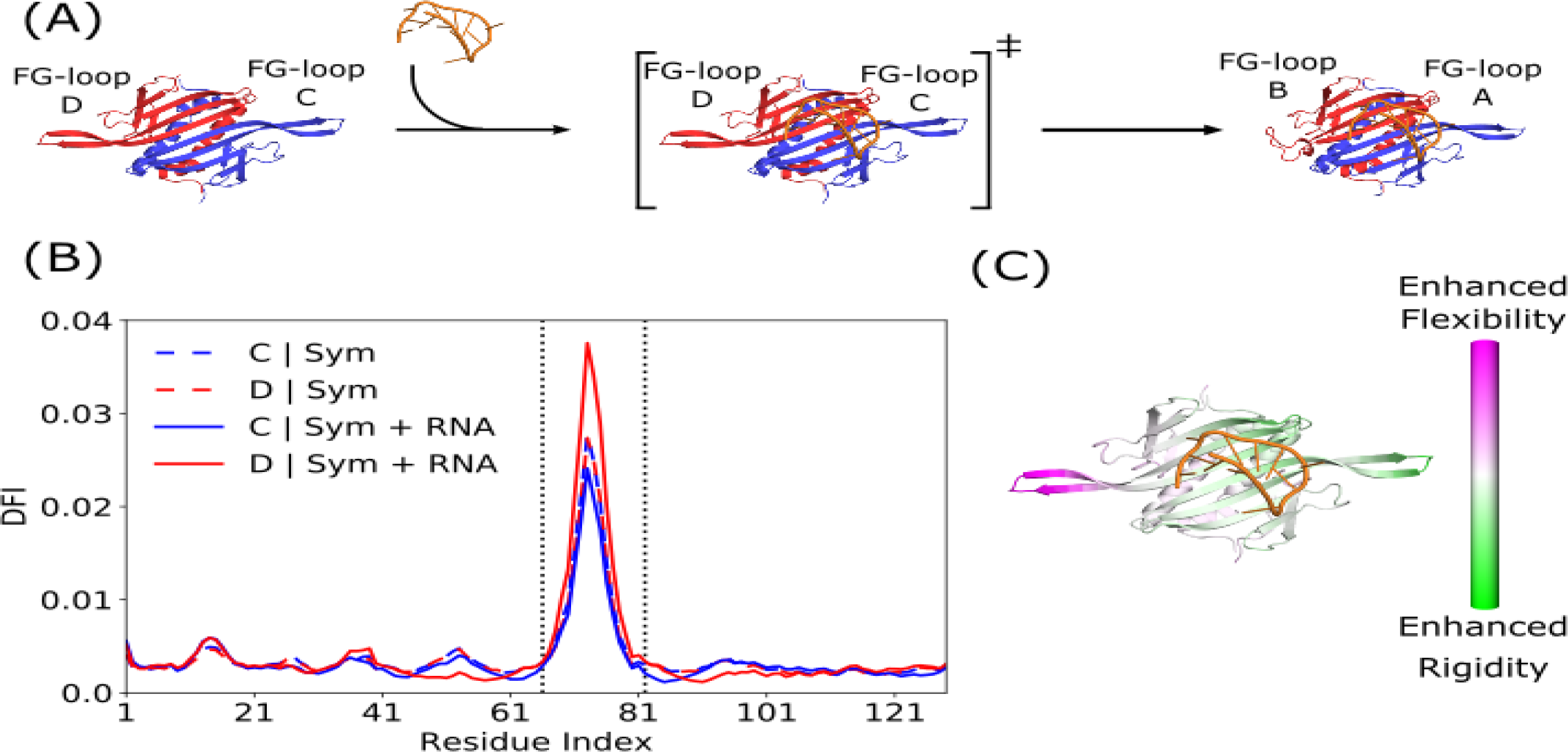
DFI profile and RNA-induced allostery in bacteriophage MS2 coat protein. (A) Cartoon diagram of the RNA-induced allostery pathway for a single coat protein dimer (chain A and C in blue; chain B and D in red). The symmetric dimer bound to RNA represents a transition state that induces a conformational change in the FG-loop furthest away from the binding site. (B) DFI profile for each of the coat protein monomers in the unbound symmetric dimer (solid lines) and the bound symmetric dimer (dashed lines). Between the dotted black lines are the FG-loop residues, which also have the highest dynamic flexibility. (C) Change in DFI from the unbound symmetric dimer to the bound symmetric dimer. RNA-binding enhances the flexibility of the FG-loop farthest away from the binding interface, indicating allosteric communication between the two domains.

### DCI analysis of wildtype coat protein dimers

The structural dynamics of the symmetric dimer are altered upon RNA-binding, as evidenced by DFI analysis. To investigate whether these alterations facilitate successful capsid assembly, we examined the DCI profiles of the wildtype unbound symmetric dimer and the wildtype bound asymmetric dimer, as shown in Fig 2. In the unbound symmetric dimer (Fig 2A), FG-loops C/D and the binding interface are strongly coupled to each other. Conversely, in the bound asymmetric dimer (Fig 2B), while FG-loops A/B are still highly coupled to the binding interface, only FG-loop A (in the flexible conformation) is highly coupled to FG-loop B (in the rigid conformation).

**Fig 2.**
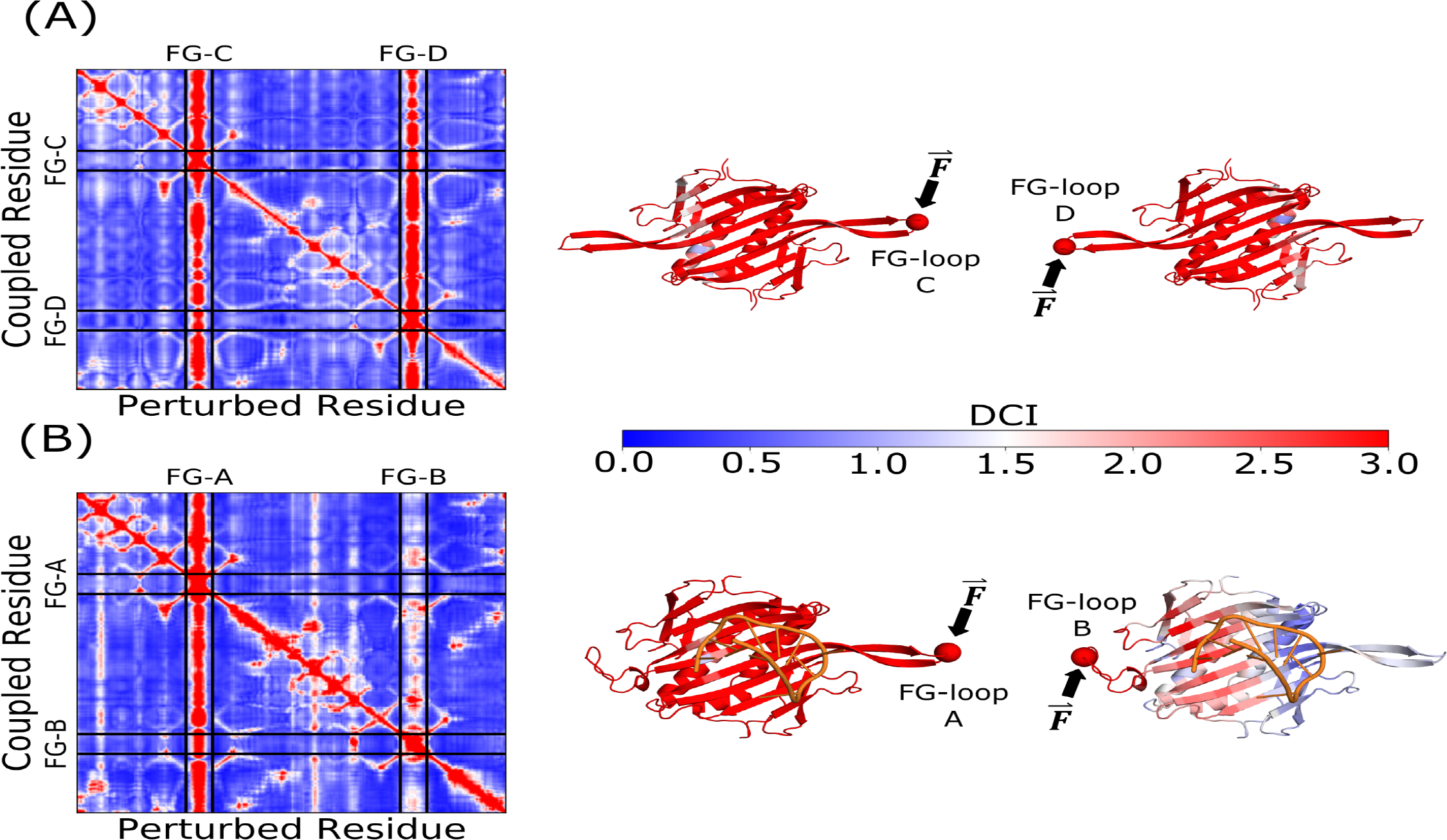
DCI profiles for the unbound symmetric and bound asymmetric wildtype MS2 coat protein dimers. (A) DCI profile for the unbound symmetric dimer. The FG-loops are symmetrically coupled, where a perturbation at one end of the dimer produces a strong response at the other end. (B) DCI profile for the bound asymmetric dimer. The symmetric coupling of the two FG-loops is broken, such that FG-loop A now drives the communication to FG-loop B. All cartoon structures are colored with DCI for perturbations at residue 74 (shown as a sphere).

Hence, the RNA-induced conformational change results in unidirectional asymmetric communication between the FG-loops, which propagates from FG-loop A to FG-loop B. These findings suggest that the flexible FG-loop A serves as the primary communicator during the initiation of capsid assembly. When the FG-loop A of an asymmetric dimer forms inter-dimer contacts with the FG-loop C/D of a symmetric dimer, it results in the long-range coupling of FG-loop A with FG-loop D/C. Following this, FG-loop D/C associates with another FG-loop A, and the system attains a minimum free energy state by bringing the two FG-loops B into contact. This is consistent with proposed assembly pathways that favor an alternating asymmetric-to-symmetric dimer association (see [31, 32]).

### Change in DCI from unbound to bound structure of wildtype symmetric, wildtype asymmetric, and mutant W82R dimers

Both the wildtype unbound symmetric dimer and the RNA-bound asymmetric dimer are crucial for correct assembly of the MS2 capsid. However, the W82R mutant dimer (PDB: 1MSC), which retains repressor activity but is deficient in capsid assembly [24, 30], poses a challenge in understanding why. To address this issue, we examined changes in dynamic coupling from the unbound to the bound state for each of the three dimers, as depicted in Fig 3 (see S1 Fig for raw DCI analysis of the mutant dimer). For the wildtype symmetric dimer, RNA-binding increases the dynamic coupling between FG-loop D and the binding interface, while decreasing the coupling between FG-loop C and both the binding interface and FG-loop D. The asymmetric shift in coupling between the two FG-loops, together with the enhanced flexibility of FG-loop D (Fig 1C), tilts the conformational ensemble of the dimer toward the asymmetric structure. However, RNA-binding to either the wildtype asymmetric dimer or the mutant dimer has a negligible effect on the coupling between FG-loops. These observations are in agreement with the previous study suggesting that the W82R mutation restricts the conformational and free energy landscape of the symmetric dimer [24, 25], possibly due to the replacement of a large hydrophobic amino acid (Trp) with a smaller hydrophilic one (Arg). Thus, we conclude that the RNA-induced change in FG-loop coupling observed in the wildtype symmetric dimer is critical for proper self-assembly of the MS2 capsid.

**Fig 3.**
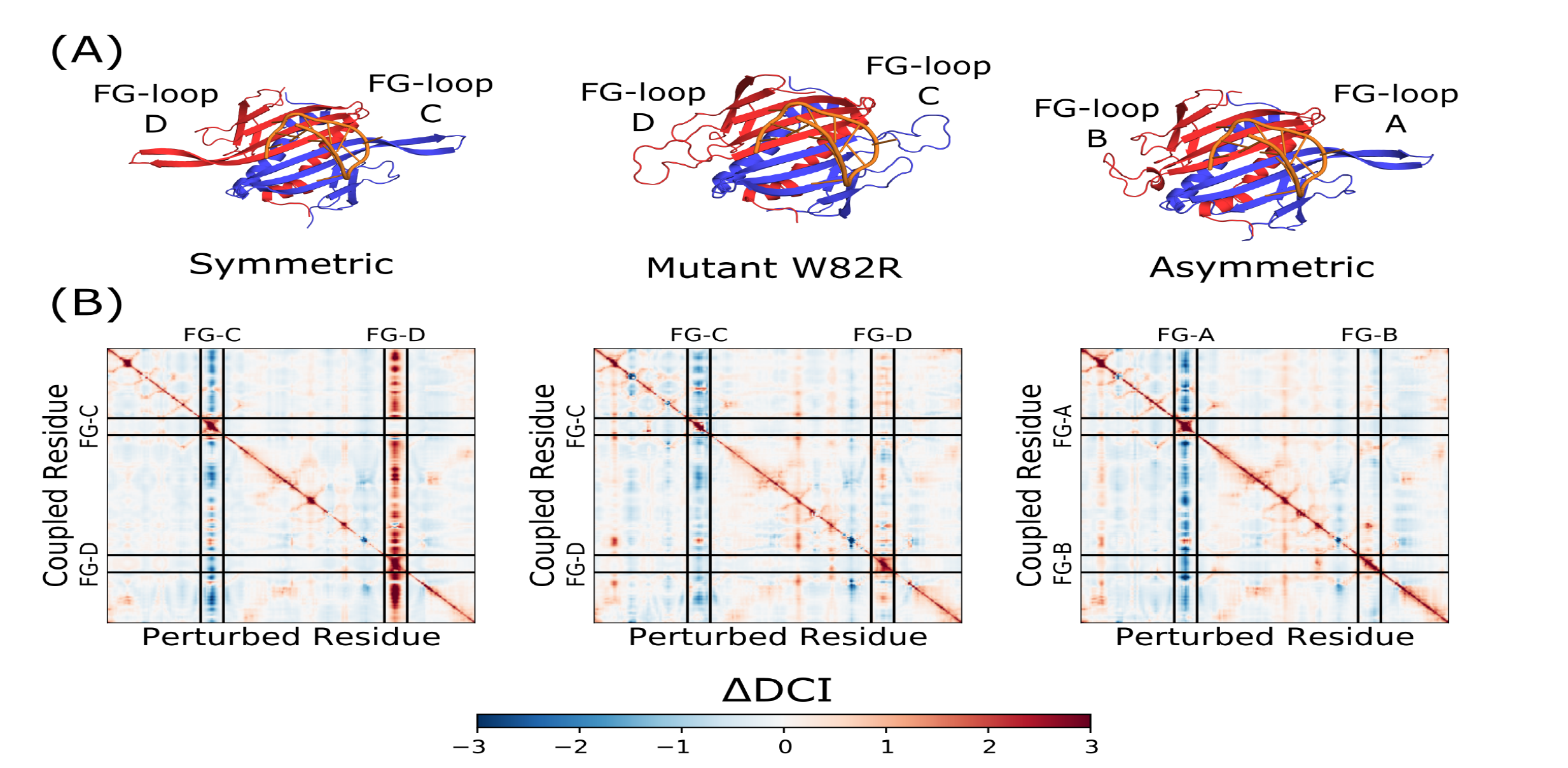
Change in DCI from unbound to bound state for three different MS2 coat protein dimers. (A) On the left and right is the wildtype symmetric and asymmetric dimers (PDB: 1ZDH), respectively, bound to RNA. In the middle is the crystal structure of the mutant W82R dimer (PDB: 1MSC) bound to RNA. In all cases, FG-loop C is in blue and FG-loop D is in red. (B) ΔDCI heat-maps for the three crystal structures shown above. Enhancement of dynamic coupling in FG-loop D is present only in the wildtype symmetric dimer, indicative of RNA-induced allostery. ΔDCI for the mutant W82R dimer and the wildtype asymmetric dimer show minimal changes in the FG-loops, indicating stable RNA-bound structures with no allosteric modulation.

### DCI analysis of MS2 capsid intermediates

The assembly pathway proposed for the bacteriophage MS2 capsid is presented in Fig 4. Although detecting protein complex intermediates can be challenging due to their short lifespan, two significant intermediates, namely the three-fold and five-fold rings, have been identified on the MS2 capsid assembly pathway via mass spectrometry [31–33]. While crescent and horseshoe structures have not been experimentally detected, theoretical evidence suggests that they are two probable structures on the pathway to the five-fold ring (see [31, 33]).

**Fig 4.**
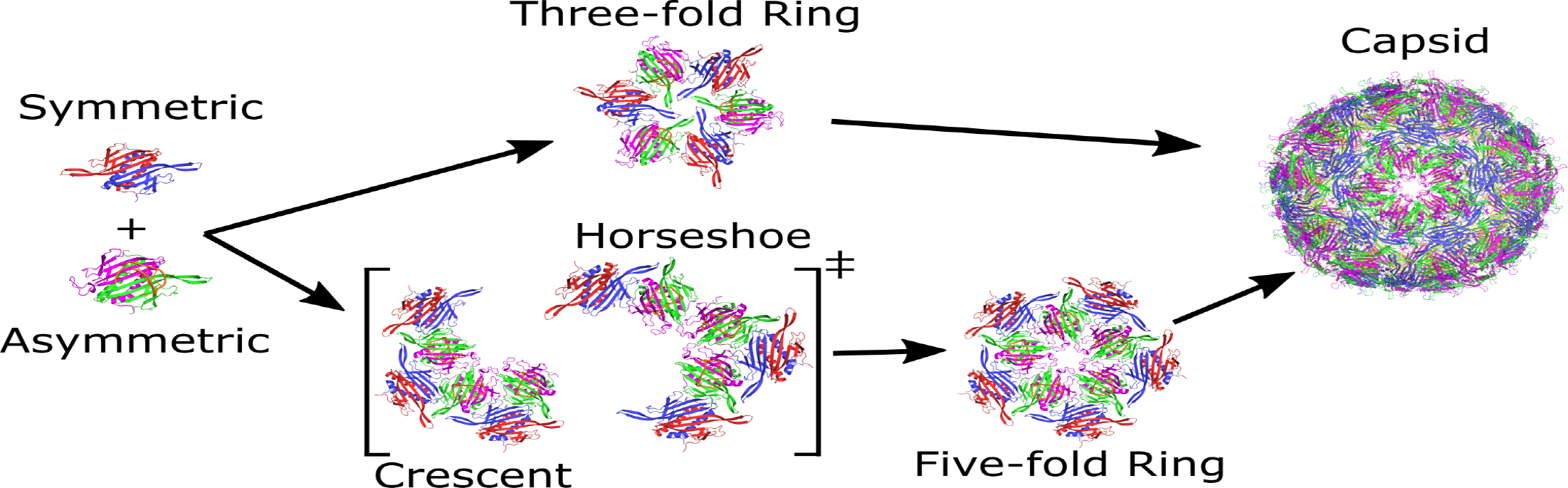
The proposed assembly pathway for bacteriophage MS2 capsid. Higher order capsid intermediates are formed by the alternating 1:1 association of asymmetric-symmetric dimers. Three-fold rings and five-fold rings form independently, with an RNA-bound asymmetric dimer acting as the nucleation site of capsid assembly. Crescent and horseshoe conformations are two possible intermediates that arise during the formation of the five-fold ring structure. In the full T=3 capsid, five three-fold rings converge at one five-fold ring.

The successful self-assembly of bio-molecules requires long-range cooperativity, which we hypothesize will be reflected in the intermediate structures along the pathway to capsid assembly. Specifically, we predict that these intermediates will exhibit high coupling in the same domains as individual coat protein dimers, namely the flexible FG-loops. To test this hypothesis, we analyzed the DCI profiles of intermediate structures perturbed at either an outer or inner FG-loop (Fig 5). We observed that perturbations at outer FG-loops, which have few to no inter-dimer contacts, exhibit high coupling that cascades throughout the entire structure (Fig 5A). Additionally, for the three-fold ring and five-fold ring, perturbations at any of the three or five outer FG-loops, respectively, result in a strong response everywhere in the structure, reflecting their rotational symmetry (see S2 Fig for more details). In contrast, perturbations at inner FG-loops, where inter-dimer contacts are fully saturated, do not propagate beyond their immediate neighbors, indicating a region of high stability at the center of the three-fold and five-fold axis.

**Fig 5.**
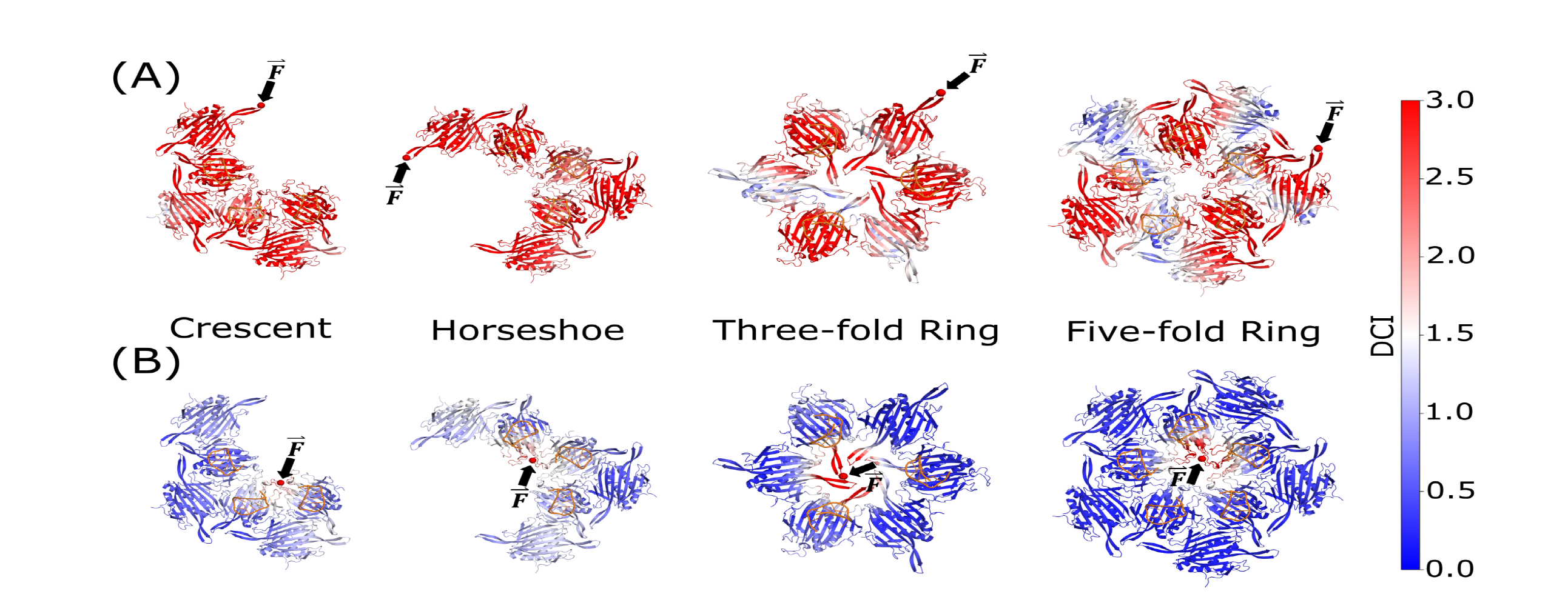
DCI for perturbations at FG-loop residue 74 in intermediate capsid structures. (A) DCI spectrum for a flexible FG-loop C/D on an outer dimer of the intermediate. These perturbations are rotationally symmetric (see S2 Fig) and highly coupled, indicating long-range communication among these FG-loops during the formation of the three-fold and five-fold axes seen in the full capsid. (B) DCI spectrum for FG-loop B on an inner dimer of the intermediate (exception: three-fold ring, which has no inner FG-loop B). In contrast to the outer FG-loops, perturbations of inner FG-loops have low dynamic coupling beyond the localized region at the center of the ring, suggesting structural stability of the FG-loop contacts at the five-fold and three-fold axes in the full capsid.

The effectiveness of the five-fold axis as a nucleation site for capsid assembly is due to the low DCI of its comprising FG-loops. These FG-loops act as a stable hinge during the formation of new inter-dimer contacts on the pathway from crescent/horseshoe to the five-fold ring, since they are weakly coupled to the rest of the structure.

Additionally, the outer FG-loops of each intermediate are flexible and highly coupled, providing the necessary mobility to guide newly-associated dimers into the correct orientation and minimizing the number of exposed flexible FG-loops. Consequently, assembly occurs spontaneously and more efficiently around the five-fold and three-fold axes, and the free energy minimization process continues until no more dimers can fit into the fully formed spherically symmetric capsid.

## Conclusion

Self-assembling viruses represent the most basic biological system that is capable of constructing itself, yet still relies on host organisms for replication. The interaction between proteins and RNA, which are ubiquitous in ssRNA viruses, is of great interest to molecular biologists and researchers seeking to develop new tools for gene therapy and drug delivery [34]. While DFI and DCI have proven effective in measuring the dynamic interactions between a small coat protein and RNA in the case of MS2, further investigations are needed to understand the mechanism in other viral species. For instance, although closely related to MS2, the FG-loops of *Levivirus* bacteriophage PP7 do not undergo conformational changes, despite similar RNA packaging signals and regulatory activity in capsid assembly [35, 36]. Moreover, the recent discovery of previously unknown variability in MS2 capsids [37] raises questions about whether the protein-RNA interactions discussed here are specific enough to support the formation of T=3 symmetry over unconventional T=4 or hybrid T=3/T=4 symmetries. Finally, the application of DFI and DCI to other viral proteins and even larger RNA-protein systems, such as ribosomes, spliceosomes, CRISPR-Cas9, and full capsids, has the potential to reveal a general allosteric mechanism among nucleoprotein complexes that can be guiding their function as well as mediate a particular directed assembly pathway, and they will be subject of our future investigations.

## Methods

### Dynamic flexibility index and dynamic coupling index

The Elastic Network Model (ENM) is a useful tool for studying protein conformational space. It represents a polymer, such as an amino acid or nucleotide sequence, as a network of nodes connected by elastic harmonic springs. ENMs have been applied to proteins in the absence of RNA, using either all-atom or coarse-grained models, depending on the specific application [38]. In recent years, the ENM has been extended and applied to RNA structures [39]. However, due to the higher flexibility of RNA compared to proteins, RNA molecules are best modeled as all-atom networks [40, 41]. For the analysis of MS2 capsid with its cognate RNA, a coarse-grained model comprised of alpha carbons was used for the viral coat protein and an all-atom model was used for the RNA-hairpin, including only the phosphate/ribose sugar backbone to ignore any sequence-specific interactions.

ENMs allow for the measurement of protein conformational dynamics at the amino acid level by simulating perturbations and measuring the response of each residue. The magnitude of fluctuation in response to random forces is directly proportional to the elastic energy of normal modes in the spring system, indicating functionally important allosteric positions [5, 6]. In this paper, two metrics, the dynamic flexibility index (DFI) and dynamic coupling index (DCI), are used to analyze structural flexibility and long-range signaling between nodes in a network. These metrics introduce Brownian forces F at each residue and calculate the response vector Δ*R* using Linear Response Theory (LRT), as shown in the equation:

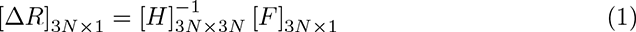

where *H* is the 3*N ×* 3*N* Hessian matrix composed of the second derivatives of the harmonic ENM potential (scaled by *R^−^*^6^ for each residue pair) with respect to the components of the 3*N* Cartesian position vectors. The Brownian forces simulate the chaotic environment within a cell, allowing for responses to be measured in a way that is most consistent with a naturally occurring system [5, 6].

From the response vectors, we construct a perturbation matrix *A* of the form:

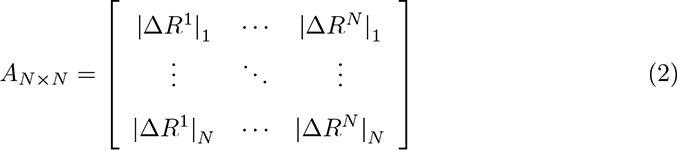

where *|*Δ*R^j^|_i_* is the magnitude of the fluctuation of residue i when residue j is perturbed. Subsequently, the DFI value of residue i is given by the net fluctuation of residue i relative to the net displacement of the entire protein:

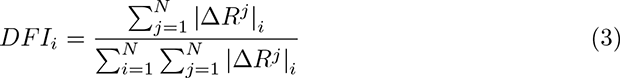

DFI is a metric that quantifies the relative flexibility of a residue, where higher values indicate more flexible positions that are functionally important (such as catalytic sites), and lower values indicate rigid positions that provide the structural integrity necessary to transmit perturbations to other residues. Computational studies using DFI have shown that point mutations at rigid positions (residues with low DFI) result in dysfunctional proteins, suggesting that these residues must be conserved across structural homologs with comparable function [5, 6]. Thus, rigid residues act as hinges (like joints/pivots) in the global motion of the protein and are sequence-specific, while flexible (high DFI) residues provide the mobility necessary for dynamical processes such as catalysis, signal transduction, and conformational changes [5, 6].

Similarly, the DCI value of residue i is given by the fluctuation of residue i in response to perturbations at a single residue j relative to the average response of residue i due to perturbations at all other residues:

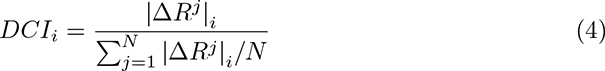

DCI is a measure of the coupling between residue i and residue j upon perturbation at residue j. If the distance between residues i and j is significant and binding of a ligand (such as hairpin RNA) increases their DCI, it suggests the presence of allosteric regulation between the sites. Previous studies on enzymes have demonstrated that certain residues distant from the active site exhibit high DCI, indicating long-range communication between functionally important residues and the allosteric sites that regulate activity [5, 6]. Thus, DFI and DCI are powerful tools for studying protein-RNA interactions and their dynamics.

Amino acid and nucleotide coordinates for the wildtype coat protein dimers (symmetric, asymmetric, crescent, horseshoe, three-fold ring, and five-fold ring) with and without RNA were generated from PDB: 1ZDH and exported via PyMOL. The mutant dimer (PDB: 1MSC) with docked wildtype RNA was generated from structure superposition on a wildtype bound asymmetric dimer using the cealign algorithm and exported via PyMOL. DFI and DCI were then calculated and plotted in Python JupyterLab using Numpy, Pandas, and Matplotlib packages.

## Supporting information

**S1 Fig.**
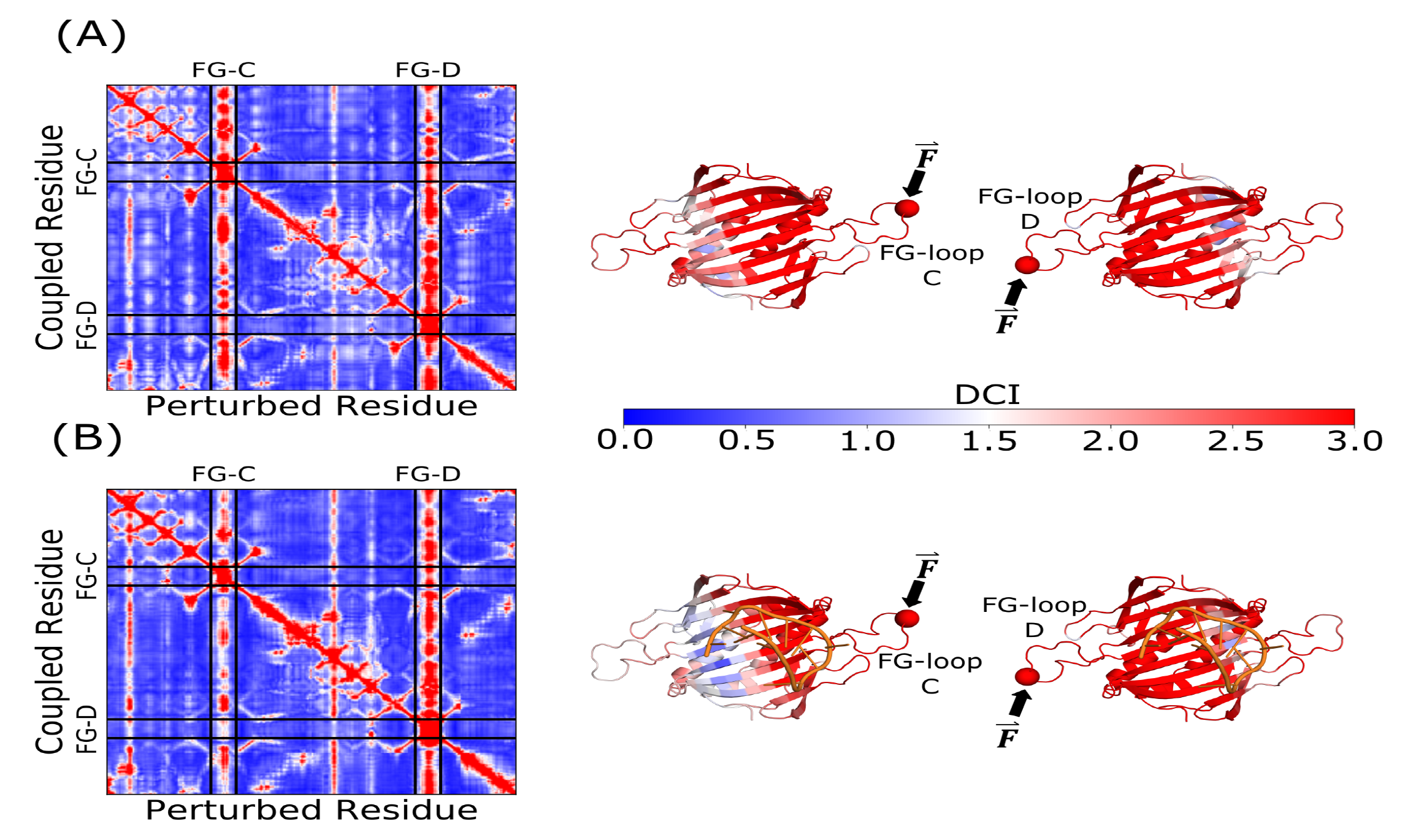
DCI profiles for the unbound and bound mutant W82R coat protein dimer that is deficient in capsid assembly. (A) DCI profile for the unbound mutant dimer. When compared with the wildtype dimer, the FG-loops are symmetrically coupled. (B) DCI profile for the bound mutant dimer. The dynamic coupling remains relatively the same as the unbound mutant dimer. All cartoon structures are colored with DCI for perturbations at residue 74 (shown as a sphere).

**S2 Fig.**
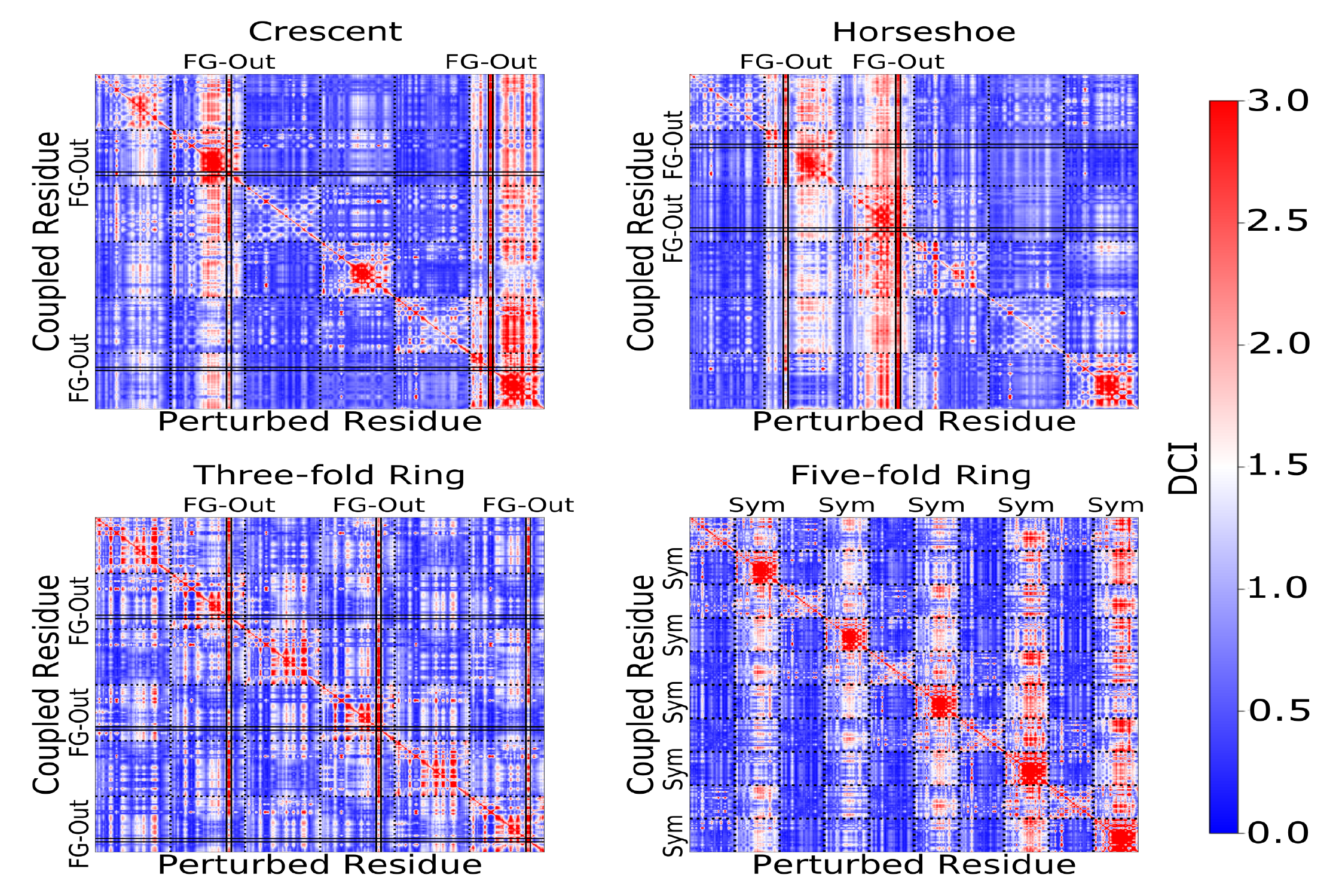
DCI profiles for the proposed intermediate capsid structures. The crescent and horseshoe structures each have two bands of high DCI that correspond to the two outer FG-loops that have no contacts to any other FG-loop. Despite the large distance between these two FG-loops, they are still highly coupled to each other. For the three-fold and five-fold rings, there are three and five bands of high DCI, respectively, corresponding to the symmetric dimers with flexible FG-loops furthest away from the center of the axis. These FG-loops are where contacts with the neighboring five-fold and three-fold rings occur. The long-range coupling ensures that proper capsid symmetry is achieved while maintaining the stability of the FG-loops that interact at these axes.

## Acknowledgments

This material is based upon work supported by the National Science Foundation under Grant No. DMR-2239518.

SBO acknowledges Gordon and Betty Moore Foundation (award number AWD00034439) and the National Institutes of Health R01GM147635-01.

## Notes

### Competing Interest Statement

The authors have declared no competing interest.

